# Machine learning and hypothesis driven optimization of bull semen cryopreservation media

**DOI:** 10.1101/2022.09.27.509504

**Authors:** Frankie Tu, Maajid Bhat, Patrick Blondin, Patrick Vincent, Mohsen Sharafi, James D Benson

## Abstract

Cryopreservation provides a critical tool for dairy herd genetics management. Due to widely varying inter- and within-bull post thaw fertility, recent research on cryoprotectant extender medium has not dramatically improved suboptimal post-thaw recovery in industry. This progress is stymied by the interactions between samples and the many components of extender media and is often compounded by industry irrelevant sample sizes. To address these challenges, here we demonstrate blank-slate optimization of bull sperm cryopreservation media by supervised machine learning. We considered two supervised learning models: artificial neural networks and Gaussian process regression (GPR). Eleven media components and initial concentrations were identified from publications in bull semen cryopreservation, and an initial 200 extender-post-thaw motility pairs were used to train and 32 extender-post-thaw motility pairs to test the machine learning algorithms. The median post-thaw motility after coupling differential evolution with GPR the increased from 52.6 ± 6.9% to 68.3 ± 6.0% at generations 7 and 17 respectively, with several media performing dramatically better than control media counterparts. This is the first study in which machine learning was used to determine the best combination of constituents to optimize bull sperm cryopreservation media, and provides a template for optimization in other cell types.

## 1. Introduction

Semen cryopreservation is a critical aspect of modern animal agriculture and the economization of animal genetic programs^1^. Moreover, semen is at the forefront of cryopreservation research, dating back to the 1600s, and was the tissue type that enabled the discovery of permeating cryoprotectants that define modern cryopreservation^2^. Freezing protocols are associated with detrimental effects on different compartments of sperm such as the acrosome, nucleus, mitochondria, axoneme and plasma membrane, consequently affecting their post-thaw fertility potential ^3^. To ameliorate this damage, sperm cryopreservation protocols use extenders to preclude both lethal and sublethal damage to sperm including preserving sperm motility, metabolic function, and fertility^3,4^. Despite large financial incentive and decades of research on semen cryopreservation, post-thaw recovery and fertility is still dramatically reduced in all species due to accumulated cellular injuries that arise throughout the freeze-thaw process^5^. The intensity of cryo-injury during freezing and thawing vary according to species and strain, but also with components and formulation of extenders, cryoprotectants used, and cooling-freezing rates^1^.

The components of the extender play a critical role in determining the post-thaw fertility of frozen-thawed sperm^6^. There are many ingredients in extenders such as membrane stabilizers (egg yolk, soybean lecithin, or milk), permeating cryoprotectants (glycerol, ethylene glycol, or dimethyl sulfoxide), buffer (TRIS or TES), sugars (glucose, lactose, raffinose, saccharose, or trehalose), salts (sodium citrate, citric acid) and antioxidants (enzymatic and non-enzymatic). In spite of decades of attempts to optimize the formulation of extenders, improvements have been limited, and as such, the majority of extenders cannot support approximately of 40% of sperm during freeze-thaw process ^1,7,8^. Conventional optimization of sperm cryopreservation has been attempted using very few changes in ingredients such as adding or removing one, two or three of extender components, but these experimental designs are generally unable to discover the effects of each individual ingredient in interaction with others. Moreover, most sperm cryopreservation studies have been fairly limited in sample size, addressing these single component differences among animal donor counts of less than 20, and typically less than 10. Because of the significant within and among animal variability in cryo-recovery, and the significant herd-to-herd, or strain to strain difference in cryo-recovery, this makes the adoption of individual studies to sperm cryopreservation at large challenging at best. In fact, this variability is a considerable challenge for sperm cryopreservation optimization. In short, there is significant variability in post-thaw recovery in sperm, even among ejaculates from the same bull^9,10^. Progress towards optimal sperm cryopreservation protocols is camouflaged by this inter- and intra-bull variability, suggesting that for any protocol to be generally optimal, a very wide selection of bulls must be made, and repeat selections from within a herd must be incorporated into the experimental design.

Sperm cryopreservation is complicated by the many interactions of the various components needed for an optimized extender of semen freezing ^11^. For example, the last 15 years of sperm cryopreservation research have been dominated by the study of antioxidants as a major component of sperm cryopreservation medium ^12,13,14^. However, since each step in the cryopreservation process and each component in cryopreservation medium has either oxidative or reductive effects, the choice and concentration of antioxidants to ensure homeostasis is challenging to generalize beyond the scope of a single experimental design. In fact, experimental evaluation of more than one or two factors (e.g. antioxidants with other extender components) at a time can be very resource intensive^15^. Therefore, to make progress towards determining optimal media components for sperm cryopreservation that accounts for the multi-factorial interactions, a new experimental strategy must be tried.

Recently machine learning was used to optimize a cell culture medium for T cells^16^ and to design an integral membrane channel rhodopsin for efficient eukaryotic expression and plasma membrane localization^17^. Similarly, a differential evolution algorithm was used to optimize cryopreservation conditions for Jurkat cells and mesenchymal stem cells^18,19^. In these experimental designs protocols or medium are optimized via information obtained through physical experiments guided by the algorithm. Machine learning has the advantage of addressing a large experimental parameter space (e.g. multiple media component concentrations) with significantly fewer total experiments than traditional factorial designs^20^. This new method can be an alternative to prevalent strategy for optimization of extender compounds with huge efficiency in process and final output^16^.

There are several possible machine learning approaches for protocol optimization. One example is differential evolution, a machine learning algorithm based on biological evolution mechanisms^21^ that can be coupled with other machine learning models via supervised learning to reduce the number of trials required. In this manuscript, we consider two supervised learning models, artificial neural networks and Gaussian process regression, to determine the optimal machine learning model for our system (see Figure 1). Artificial neural network models require no prior model knowledge or intuition regarding the relationship between the inputs and experimental results during the model training process^22, 23^. This allows the artificial neural network to approximate a wide range of different models ^24^. Gaussian process regression models are probabilistic and have been used to analyze and optimize media composition for lipid productivity^25^. Unlike the artificial neural network, Gaussian process regression incorporates error estimates in its model predictions, and accounts for the error in experimental results^25^.

**Figure 1:**
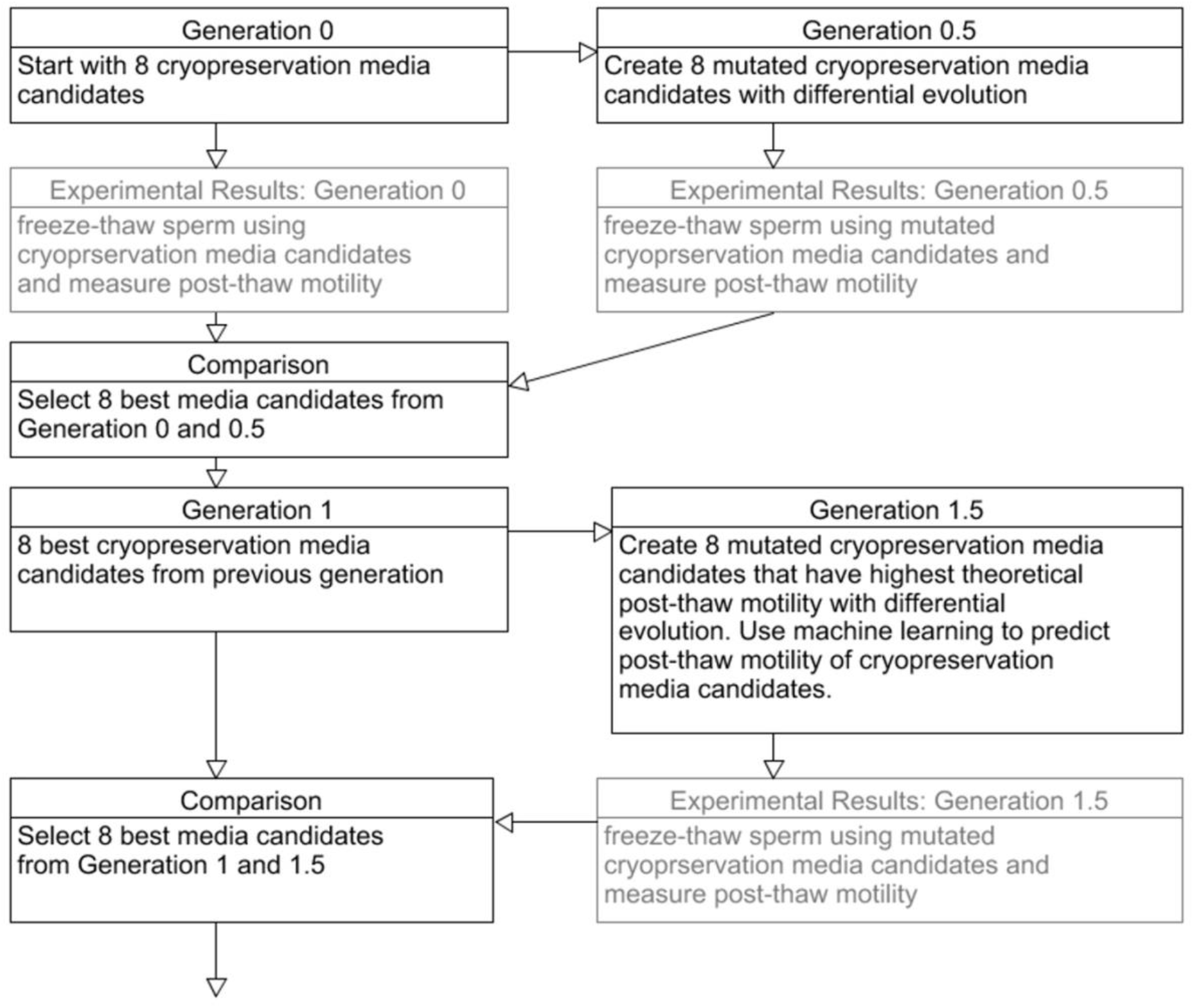
Flow chart of experimental design.

A broad range of ingredients with preservative characteristics have been used in semen extender during cryopreservation. For bull sperm, egg yolk^26,27^ or milk^28^ are the major extracellular components to protect the plasma membrane against cryopreservation damage through the dynamic substitution of membrane phospholipids that prevent or ameliorate membrane phospholipid loss^29,30^. The first discovered permeating cryoprotectant, glycerol, is another key component of most bull sperm extender media, though some reports of the success of the permeating cryoprotectant ethylene glycol have been published^31^. Both of these chemicals act as the main permeating cryoprotective agent (CPA) to support dehydration at lower temperatures, reduce intracellular ice formation, and thus increase survival during cryopreservation^14^. Sugars such as fructose, sucrose, and trehalose are another common ingredient of extender media that provide some protection during cryopreservation, though typically sugars are not able to diffuse across the plasma membrane, instead creating an osmotic pressure in the media that induces cell dehydration, a lower incidence of intracellular ice formation, and an increased likelihood of vitrification in the unfrozen space surrounding sperm at low temperatures^32^. Sucrose and trehalose are considered standard extracellular CPA and are associated with the promotion of extracellular glass during cooling, but also membrane stabilization and ice recrystallization inhibition^33,34^. Fructose is generally an energy source for sperm, but it also facilitates adjusting the osmotic pressure of the extender and acts as a cell protectant^35^. The combination of sugar and tris (hydroxyl-methyl) aminomethane (Tris) buffer has a great impact on the success of the cryopreservation medium^36^.

During cryopreservation, sperm are subjected to many biochemical, mechanical, and ultrastructural stresses, causing detrimental effects on post-thaw parameters and fertility potential^37^. Reactive oxygen species (ROS) are a major source of this damage and many studies have hypothesized that antioxidants would be able to support sperm by counteracting ROS during the cryopreservation process. While there are myriad choices for antioxidant classes and species, here we focus on two: melatonin, a secretory product of the pineal gland that serves as an effective antioxidant and eliminates ROS during cryopreservation^38^, and nerve growth factor (NGF), an antioxidant that supports cellular signal transduction and improves capacitation and acrosome protection which is one of the critical characteristics of sperm during fertilization which needs to be protected during freeze-thaw^39,40^.

In this manuscript we propose to identify an optimal bull sperm cryopreservation medium from a set of components listed in Table 1. To identify the optimal concentration of each component in this medium, we employ an iterative feedback loop informed by supervised machine learning and experimentation on semen from a large herd of production bulls. This process was repeated over 11 generations of media designs and resulted in significant improvement in post thaw recovery against industry standard medium in a large sample group of bulls.

**Table 1:** Media components used in iterative design and their purpose. Each component concentration is varied during each trial according to predictions from the simulated annealing algorithm.

| ***Design Variable*** | ***Purpose*** | ***Initial Concentration*** |
| --- | --- | --- |
| Tris | Base media | 0-50 % |
| Egg yolk | Membrane stabilizer | 0-20 % |
| Milk | Membrane stabilizer | 0-15 % |
| Fructose | Energy source | 0-1.25 % |
| Glycerol | Permeating cryoprotectant | 0-7% |
| Ethylene Glycol | Permeating cryoprotectant | 0-6.75% |
| Trehalose | Non-Permeating cryoprotectant | 0-100 mM |
| Cholesterol loaded cyclodextrin | Membrane stabilizer | 0.5-1 mg/ml |
| Glutathione | Enzymatic antioxidant property | 0.5 mM |
| Melatonin | Antioxidant | 2-3 mM |
| Nerve Growth Factor | Growth factor | 0-0.5mg/ml |

## 2. Results

### 2.1 Optimal Theoretical Model

To determine the optimal model, 200 extender-post-thaw motility pairs were used to train and 32 extender-post-thaw motility pairs test the machine learning algorithms. Cross validation produced an average mean squared error of 525 and 576 for the artificial neural networks and Gaussian process regression respectively. Post-thaw motility predictions by the artificial neural network and Gaussian process regression had mean squared error 275 and 883 respectively. Overall, the artificial neural networks had less prediction error then the Gaussian process regression.

The median post-thaw motility at generation 6 for the artificial neural networks predictions, Gaussian process regression predictions, and experimental results were 43.6 ± 2.9%, 36.0 ± 20.7%, and 47.4 ± 16.5% (median ± SD) respectively. The Kruskal-Wallis test indicated that there was no significant difference between artificial neural network predictions, Gaussian process regression predictions, and experimental results (χ^2^ = 3.90, df = 2, p = 0.14). Comparing pair wise predictions with the Dunn test showed that artificial neural networks and Gaussian process regression predictions were not significantly different (p=0.196). The Fligner-Killeen test showed that experimental results and the artificial neural network predictions were heteroscedastic (χ^2^ = 9.23, df = 1, p = 0.002) and homoscedastic with the Gaussian process regression (χ^2^ = 1.20, df = 1, p = 0.27; Figure 2).

**Figure 2:**
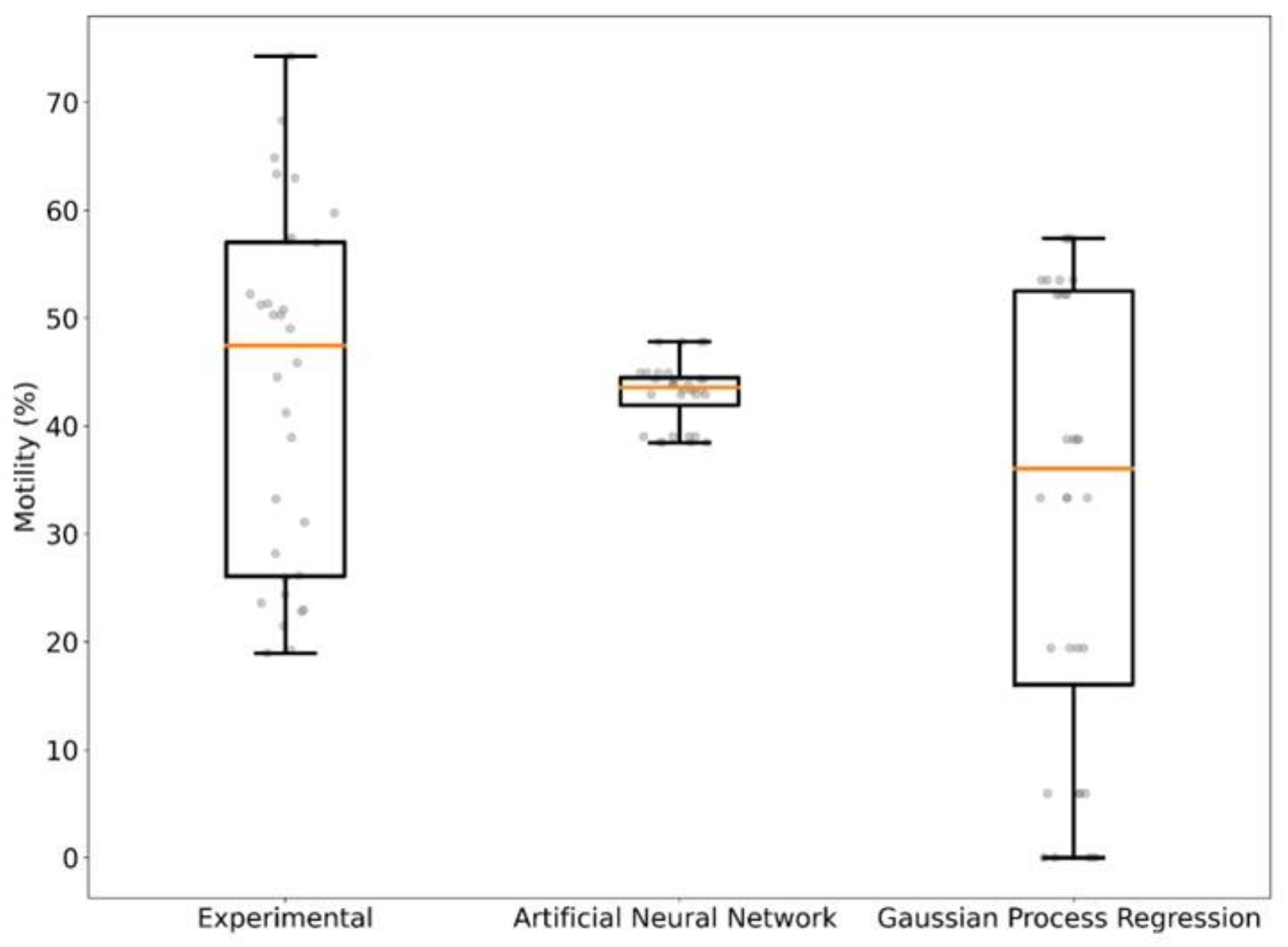
Experimental results and post-thaw motility predictions for Generation 6, where artificial neural network and Gaussian process regression models were trained using results from generation 1—5.

### 2.2 Optimum extender composition

The median post-thaw motility from generation 1 to 6 increased from 25.5 ± 20.3% to 47.4 ± 16.5% (median ± SD) based solely on differential evolution. After coupling differential evolution with GPR the median post-thaw motility increased from 52.6 ± 6.9% at generation 7 to 66.7 ± 3.1% at generation 17 (median ± SD) using total motility as the optimization metric. The median post-thaw motility increased from 50.0 ± 14.0% at generation 18 to 68.3 ± 6.0% at generation 21 (median ± SD) using relative total motility as the optimization metric (Figure 3).

**Figure 3:**
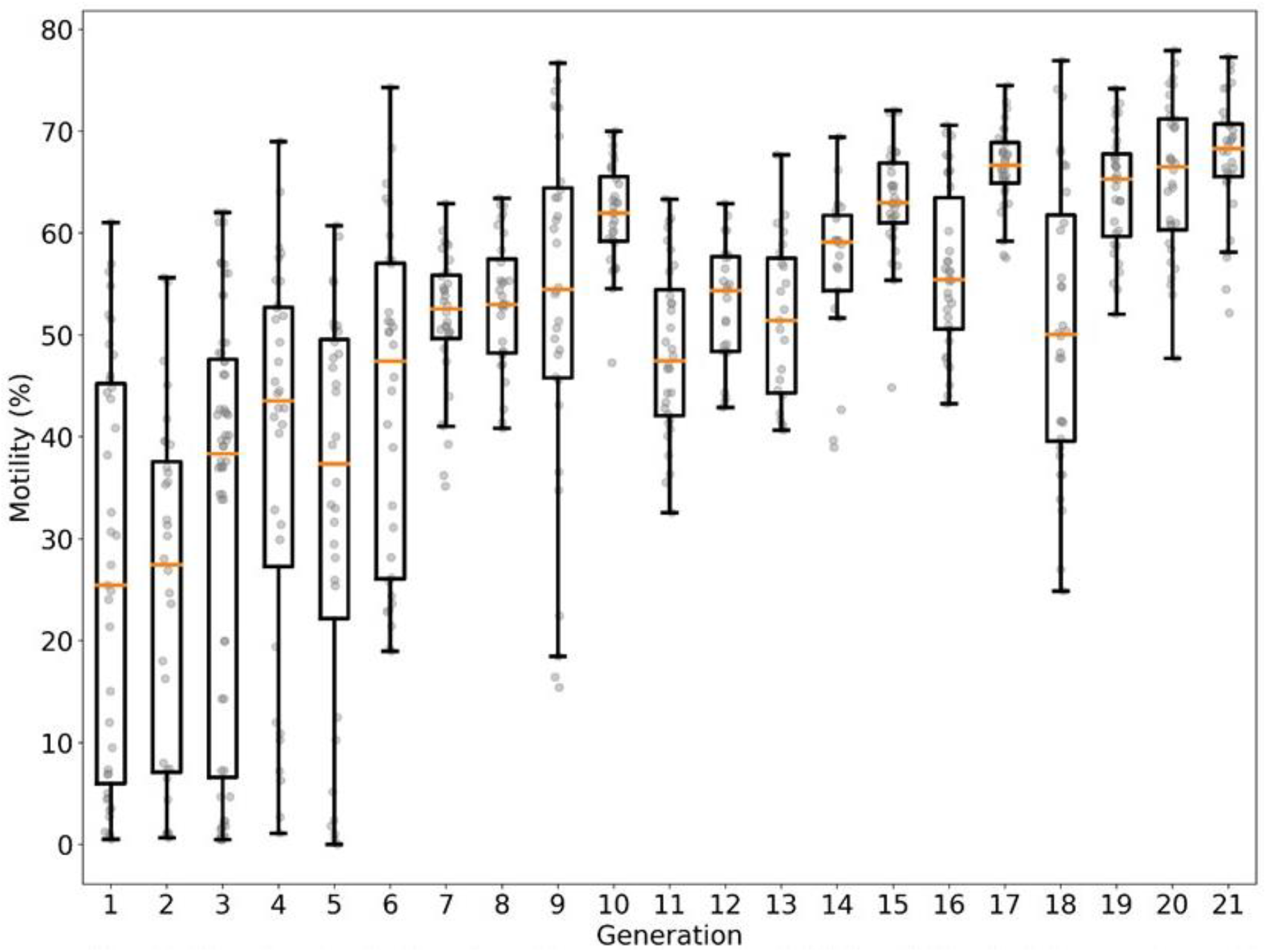
Generational results for total motility metric (generation 1-14, 16, and 17) and relative post thaw motility metric (generation 18-21). Generation 1 to 17, 18, to 21 correspond to extender compositions generated by differential evolution. Generation 15 corresponds to re-testing of the top performing extenders from generation 1 to 14.

The median post-thaw motility for the total motility metric at generations 15, 16, and 17 were 63.0%, 55.4%, and 66.7% respectively. At generation 17, median post-thaw motility for the control medium was 70.5%. There was no difference in motility between the total of generation 17 extenders and commercial medium (χ^2^ = 1.58, df = 1, p = 0.21). The median post-thaw motility for the relative total motility metric at generations 19, 20, and 21 were 63.3%, 66.5%, and 68.3%, respectively. At generation 21, median post-thaw motility for the control medium was 65.3%. The median post-thaw motility for all but one extender in generation 21 exceeded the control medium. However, there was no difference in motility between the total of generation 21 extenders and commercial medium (χ^2^ = 2.13, df = 1, p = 0.14). The convergence testing between generations 19, 20, and 21 indicated that there were no differences between the three generations and the algorithm was approaching a set optimal media composition (χ^2^ = 4.31, df = 2, p = 0.12). We did not observe any difference in post-thaw between total post-thaw motility or relative post-thaw motility optimization metric (χ^2^ = 1.73, df = 1, p = 0.18. (Figure 4).

**Figure 4:**
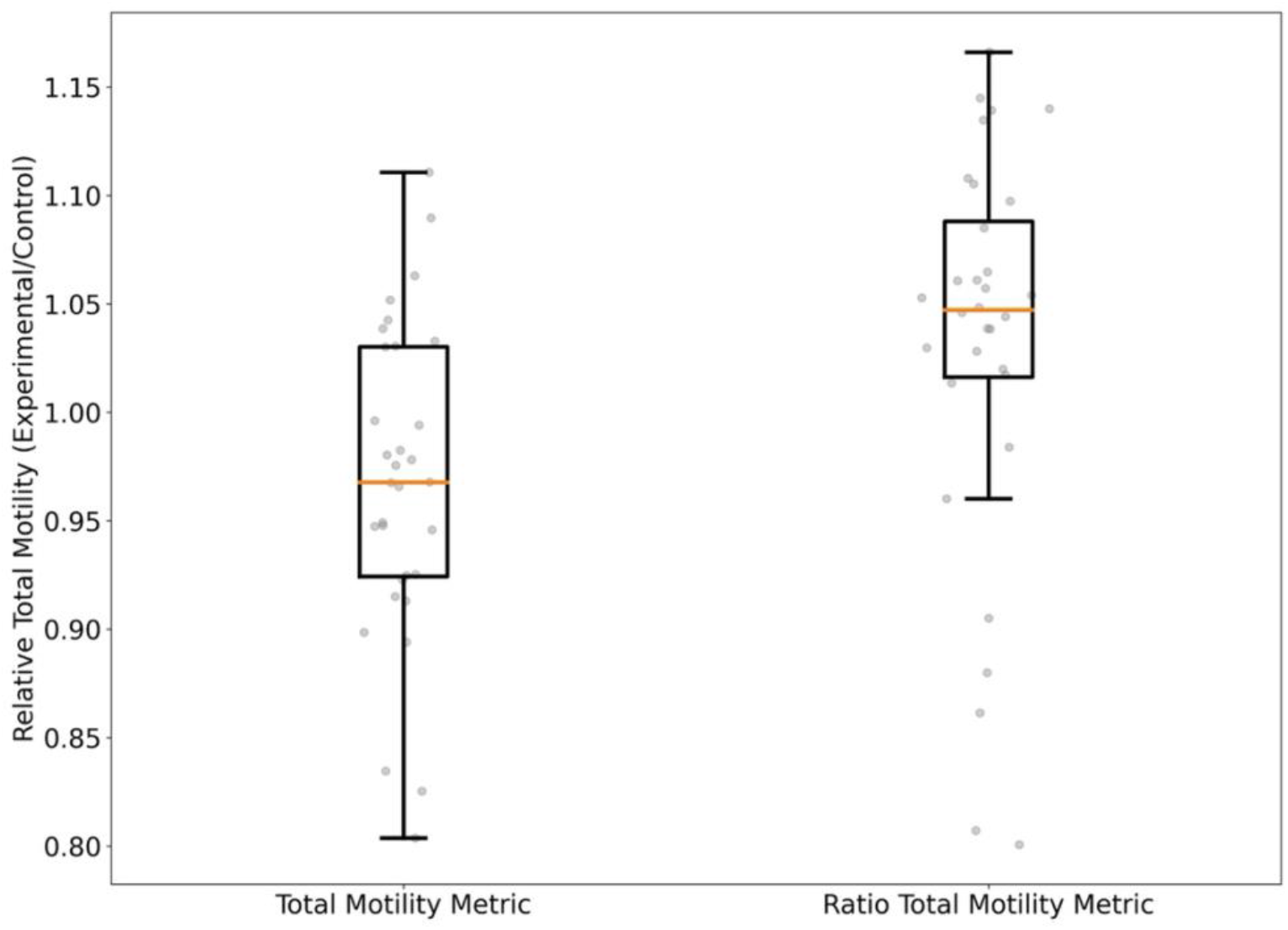
Direct comparison of media generated by differential evolution to commercial medium for media at generation 17 using the total motility metric and at generation 21 using the relative total motility metric.

### 2.3 Validation of top media designs

A set of five out of 196 algorithm driven extenders were chosen to compare with the standard control egg-yolk based extender. The criteria for choosing the 5 extenders were relative post-thaw motility (see Figure 5). The mean post-thaw motility and progressive motility were 31.9 ± 1.3% and 23.2 ± 1.1%, respectively, for the top five extenders, compared with 25.2 ± 1.1% and 16.8 ± 1.1%, respectively, for the corresponding controls. This corresponds to a 26.6% and 37.6% improvement in post-thaw motility and progressive motility, respectively. Table 2 shows the components of the top extenders and concentrations of each individual component. The results for post-thaw motion characteristics and flow cytometry parameters of top extender and standard control extender are presented in Table 3. We used a stepwise approach to identify the top algorithm-driven extenders by comparing the various motion characteristics and flow cytometry parameters in post-thaw semen in the top algorithm driven extender and the commercial extender as control. Total motility in the top extender varied between 57.4% to 63.4% which is comparable to the amount of control extender with 60.5% value. There is a similar trend for progressive motility so that the average of post-thaw progressive motility was equivalent to the average of control post-thaw progressive motility (Table 3). While all the kinetic parameters (VAP, VCL, ALH and BCF) had similar value compared to the control extender, LIN as important fertility associated metric was significantly higher in the algorithm driven extenders (Table 3). There was no significant difference in terms of percentage of mitochondrial activity, membrane functionality and acrosome integrity within and between 5 top extenders and standard control extender (Table 3).

**Figure 5.**
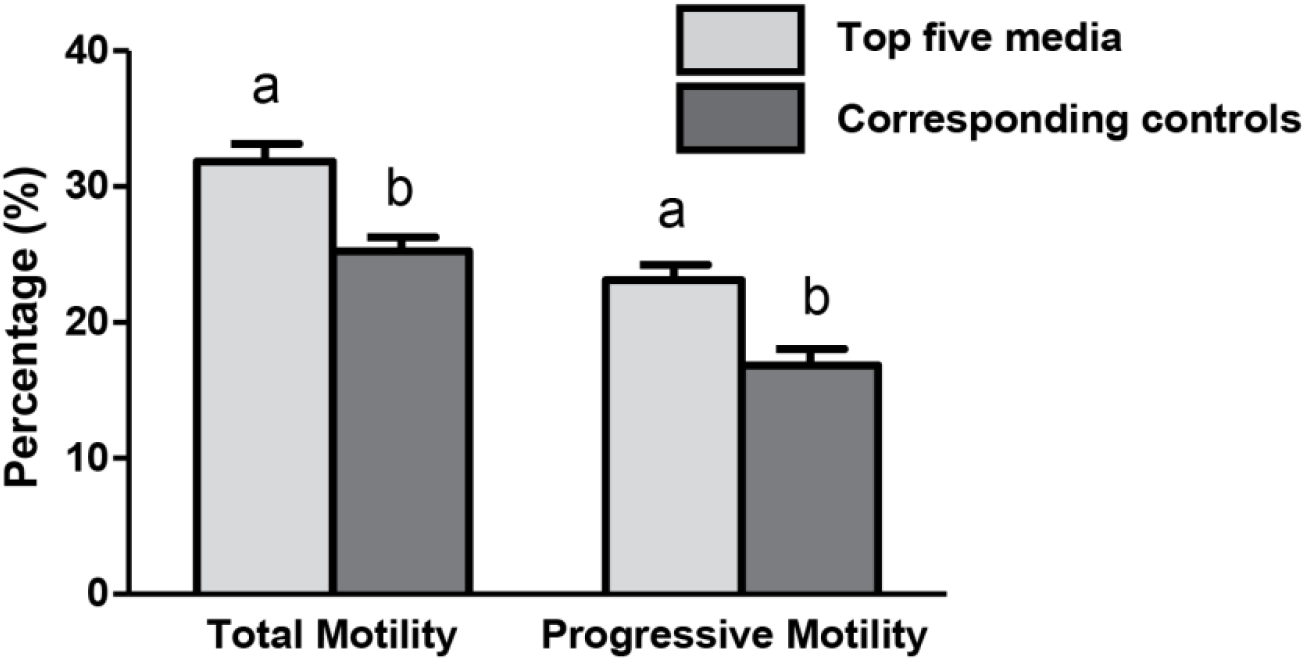
Average of total and progressive motilities between top five media recipes and their corresponding values for control extenders (split sample) in four bulls (*n*=20). Error bars indicate SEM. A significantly higher percentage of total and progressive motilities were observed in these selected extenders compared to standard control extenders (p<0.05; different letters in each category). Total motility and progressive motility were measured by computer assisted semen analysis (CASA).

**Table 2.** The components and concentrations of the five top extenders driven by algorithm design.

| Extender | Water (mL) | Tris (mL) | Egg Yolk (mL) | Milk (mL) | Fructose (mg) | Trehalose (mg) | CLC (mg) | Glutathione (μL) | Melatonin (μL) | NGF (ng) | Glycerol (mL) | Ethylene Glycol (mL) |
| --- | --- | --- | --- | --- | --- | --- | --- | --- | --- | --- | --- | --- |
| Extender 1 | 1.17 | 6.31 | 1.88 | .07 | 44.8 | 162.3 | 6.9 | 6.1 | 8.7 | 245.2 | 0.61 | 0.0 |
| Extender 2 | 1.22 | 7.28 | 1.87 | 0.05 | 0.09 | 138.3 | 6.7 | 5.6 | 10.7 | 213.2 | 0.61 | 0.0 |
| Extender 3 | 0.57 | 6.68 | 2 | 0.17 | 46.1 | 147.0 | 5.1 | 3.1 | 8.7 | 211.2 | 0.65 | 0.07 |
| Extender 4 | 0.16 | 7.3 | 1.87 | 0.3 | 0.43 | 100.0 | 6.3 | 5.1 | 9.4 | 209.2 | 0.64 | 0.1 |
| Extender 5 | 1.16 | 5.8 | 2 | 0.01 | 59.8 | 142.3 | 5.1 | 5.0 | 9.2 | 230.2 | 0.61 | 0 |

**Table 3.** Post-thaw motion and flow cytometry parameters of bull sperm in 5 top algorithm driven extenders

| Treatment | TM | PM | VAP | VCL | LIN | ALH | BCF | MF | MA | AR |
| --- | --- | --- | --- | --- | --- | --- | --- | --- | --- | --- |
| Extender 1 | 58.9 | 38.5 | 75.8 | 148.4 | 56.7 | 3.1 | 30.2 | 63.4 | 55.6 | 11.4 |
| Extender 2 | 60.2 | 40.1 | 73.2 | 145.2 | 63.5 | 3.5 | 28.4 | 60.4 | 57.3 | 9.4 |
| Extender 3 | 57.4 | 35.4 | 76.1 | 154.8 | 58.9 | 2.9 | 33.1 | 58.4 | 59.3 | 14.3 |
| Extender 4 | 63.4 | 43.2 | 73.5 | 139.5 | 64.3 | 3.2 | 27.4 | 64.3 | 54.2 | 12.9 |
| Extender 5 | 60.1 | 39.5 | 77.9 | 142.1 | 60.1 | 3.4 | 35.6 | 66.1 | 58.3 | 13.56 |
| Control | 60.5 | 41.2 | 76.6 | 139.6 | 53.6 | 3.3 | 32.4 | 63.4 | 55.6 | 11.4 |
| SEM | 2.69 | 1.98 | 3.24 | 3.19 | 2.48 | 1.18 | 1.56 | 2.5 | 1.89 | 1.46 |

There were no significant differences between top dot and control extenders. Kinetic parameters were measured by computer assisted semen analysis (CASA). TM, total motility; PM, progressive motility; VSL, progressive velocity; VCL, curvilinear velocity; LIN, linearity; BCF, beat cross frequency; VAP, average patch velocity; ALH, amplitude of lateral head displacement. Cellular parameters were measured by flow cytometer. MF, sperm with membrane functionality; MA, sperm with mitochondria activity; AR, sperm with acrosome reaction,

## 3. Discussion

Extender components play a critical role in protecting sperm during cryopreservation, and each component has a specific role as a stabilizer, cryoprotectant agent, energy provider, or antioxidant. Extenders are composed of a number of these ingredients and there are multiple interactions between each individual component. Conventional methods of extender development are not able to discover the effects of each individual ingredient in interaction with others and often are only able to evaluate one or two factors at a time. These complications are compounded by the high within- and among-bull post thaw recovery from sample to sample. Therefore, our approach allows an experimental exploration of these interactions to determine optimal freezing media. In the current study, we used machine learning to optimize the composition of bull semen freezing media, with the hope that it would provide a tool robust against the widely varying within and among-bull post-thaw fertility. This is a new strategy in bull semen cryopreservation.

The first objective of this study was to evaluate the post-thaw motility between two machine learning models, artificial neural networks and Gaussian process regression. An artificial neural network is generic in structure and possesses the ability to learn from historical data as well as with relatively less data compared to Gaussian process regression ^22^. Bas and Boyasi ^23^ used similar modeling techniques for enzyme kinetics, and found that artificial neural networks are better than response surface methodology for both data fitting and estimation capabilities ^23^. We observed the same result within our study. The artificial neural network produced a smaller mean squared error than the Gaussian process regression model. However, the Kruskal-Wallis test indicated that there was no significant difference between the experiential results and either of our two models. This may indicate that using mean squared error to as a metric for selecting the optimal theoretical model is insufficient.

Although the mean squared error showed that the artificial neural network model is superior to the Gaussian process regression, it may not be the optimal model. In fact, there were large differences in experimental predictions. The artificial neural network model predicted a small variation in the post-thaw motility compared to the experiential data where there is large variation in post-thaw motility. This may indicate that artificial neural network models are more suitable for the cryopreservation of homogeneous samples like cultured Jurkat cells or mesenchymal stem cells^18^. However, sperm samples are much more heterogeneous, with random sampling, genetic difference, and seasonal fluctuation producing samples that are highly variable. As such, we observed that the Gaussian process regression produced variance that was more comparable with experimental results. As a model, the Gaussian process regression model accurately predicted experimental post-thaw motility and captured the sample variance, whereas the artificial neural network was only able to match post-thaw motility predictions.

The second objective of this study was to optimize the formulation of ingredients in extender for improved post-thaw motility in bull sperm. To do this, we combined Gaussian process regression and differential evolution to optimize the extender components to producing media that was comparable to our commercial control extender. We needed 21 generation to create extenders that was comparable to commercial that have been optimized over 60 years. Our approach also required significantly fewer trials compared to a simple factorial approach, which, with three levels (e.g. zero, low, high) for each factor (a crude estimate of real protocols), would have needed nearly 200000 treatment combinations, and due to the variability among and even within bull semen, the number of experiments for this many treatments would be impossible.

Differential evolution is a parallel direct search method based on biological evolution mechanisms^21^. Pollock et al.^18^ were the first to apply differential evolution to optimize media composition and cooling rate for the cryopreservation of cultured Jurkat (white blood) cells and mesenchymal stem cells^18^. We found similar results when applying these techniques to bull sperm cryopreservation. This iterative approach has several benefits compared with conventional experimental factorial approaches, including adaptation to new samples, the ability to add/subtract ingredients easily and specify for low cryotolerance. Thus, we expect that this modeling-informed experimental design can be applied to other species’ sperm, and the cryopreservation of other nonhomogeneous cells and tissues. In the current study, mean post-thaw motility from generations more often increased each generation with the total motility metric. With the relative total motility metric, we see that generation 19 - 21 the algorithm post-thaw motility began to plateau. The final generation for each optimization metric (Figure 3), we observed that differential evolution using relative total motility produced more media that were on-par or better then control.

During cryopreservation, extender components and the their interactions dramatically affect sperm survival^41^. Motility of sperm is the main frequent quality control parameter for selecting sperm with the highest fertility potential^41^. There is strong positive correlation between motility and fertility. Therefore, we selected motility as the main criteria to select the best extenders in each generation. The post-thaw motility of sperm in the latest version of designed extender was comparable to the commercial control extender which is in the regular use for the artificial insemination. To validate motility data, we performed a flow cytometric experiment and there was a logical relationship between the mitochondrial activity, apoptosis, and acrosome integrity with the post-thaw motility observed in the previous step.

In the current study, extender formulations were parameterized using a number of components including buffer, sugars, CPAs, antioxidants, and membrane stabilizers. These parameters defined the extender recipes that were tested experimentally. The output of the experiment (total motility) was the input into the algorithm that prescribed the next set of extender components. This process was repeated sequentially until a stopping condition was observed. On average our post thaw motility increased 0.5% per trial and some trials had media exceeding post thaw motility from standard protocols. Based on the post-thaw recovery in the best media, we repeated the testing of five final media obtained from our algorithmic approach with comparable or superior post-thaw recovery compared to the control standard extender. Unfortunately, the gains seen in previous trials during the main experiment where recovery was well above control medium were not seen as dramatically in our follow up experiment. However, an improvement in kinetic parameters specially LIN was observed in our top media. This improvement is significant as it has certain impact on field fertility. The algorithm driven extenders have had variety of compounds that can stabilize the membrane integrity of sperm as well as osmotic tensions during freeze-thaw. Cholesterol loaded cyclodextrin and trehalose were two major compounds of our extenders that are thought to be membrane stabilizers facilitating osmotic resistant against damage.

During the optimization process the differential evolution algorithm did eliminate some variables and we manually removed others as new research emerged. There are many studies that compare glycerol and ethylene glycol as CPAs^42,43,44,45^. Recently, it was shown that may be possible to replace glycerol with ethylene glycol without effecting sperm quality or cryosurvival^42^. These studies looked at individual effects of glycerol and ethylene glycol on post-thaw parameters, as ethylene glycol and glycerol both perform similar roles as a permeating CPA. We initially hypothesized that that an extender with both ethylene glycol and glycerol may increase post-thaw motility. However, this was not the case. We found that extenders that contained ethylene glycol almost universally had lower post-thaw motility then those with glycerol only. The differential evolution algorithm often generated extenders that had zero ethylene glycol during the optimization process. At generation 21, we found that the extenders with the highest post-thaw motility contained no ethylene glycol. The differential evolution algorithm also eliminated fructose in some trials. We know fructose is an integral component to extender recipes^3^, and it would be uncontroversial to state that removing it could have been detrimental. However, we observed the opposite: extenders without fructose had good post-thaw motility. It is unknown if the positive result was merely caused by interactions between components in the extender. We leave the exploration of the component interactions for future work. The differential evolution algorithm included growth factor for 18 generations, however work by Kowser et. al.^46^ indicated that growth factor may be detrimental to post-thaw motility. Thus, to explore this effect and the hypothesis that growth factor may be detrimental, we manually removed growth factor for generation 19. We found that post-thaw motility increased after removing the growth factor. Note, however, if this was not the case, we could have continued and used the original recipe with growth factor included, without any loss or setback. As all additional data could be used to further train the Gaussian process regression. This shows that our optimization approach makes it possible to incorporate new information and explore ongoing hypotheses during the whole optimization process without any negative impact (e.g. wasted time or resources).

We were able to replicate some industry standards recipes after 21 generations (Table 2). Glycerol is a common CPA used many extenders. The standard glycerol amount is 6 % (v/v)^47^. The average glycerol amount we arrived at was approximately 6% (v/v). Typical tris-based extenders have 20% (v/v) egg yolk, our extenders only needed slightly less egg yolk. Standard fructose concentration is about 55.5 mM^35^, and our new extenders that had fructose only needed either slightly less than the standard (50 mM) or very minimal amounts. This reduction may simply be due to the number of extender components that we explored, which reduced the amount of egg yolk and fructose that was needed. Our approached allowed us to replicate industry standards as well as reduce the amount or concentration of other necessary components. Therefore, we have demonstrated that this approach is a feasible methodology to verify that current extenders are optimal, to reduce the cost of manufacturing extenders, and to develop complex cryopreservation medium for cells and tissues where there is not a mature cryopreservation medium.

## 4. Conclusions

In this study, the Gaussian process regression model was optimized for predicting post-thaw motility of frozen extender in extenders generated by differential evolution. These approaches are the first steps towards improving post-thaw functionality and still need to be defined based on multiparameter fertility metrics such as a combination of motion and flow cytometric parameters. The post-thaw results obtained by this approach were comparable to commercial and industry standards, however only a field fertility trial will provide complete evidence of the true fertility of each sample as a function of preservation medium.

## 5. Materials and methods

### 5.1 Chemicals and reagents

All chemicals used in current study are provided by Sigma (St. Louis, MO, USA), and Merck (Darmstadt, Germany) Company, unless otherwise indicated. All animal experimental procedures were approved by the University of Saskatchewan University Animal Care Committee (UAP 002CatA2018), and ARRIVE guidelines have been followed in the methods.

### 5.2 Bulls and semen collection

Ejaculates were collected using an artificial vagina from 68 Holstein bulls, aged between 15 and 30 months, regularly used for breeding purpose at Semex (Quebec, Canada). Semen samples were kept in a water bath at 33 °C while preliminary analysis of fresh semen was performed to measure the concentration, motility, and morphology. Samples that did not meet minimum quality assessments (namely greater than 10^9^ sperm/mL, motility greater than 70%, and less than 15% abnormal morphology) were excluded from the study. Semen samples used in the study were collected as part of the regular production schedule from a production herd, and as such were a random selection of the greater production herd (~200 bulls) on site.

### 5.3 Extender preparation and cryopreservation

A standard control tris-based extender was prepared by dissolving 200 mmol/L tris (hydroxymethyl-aminomethane), 66.7 mmol/L citric acid and 55.5 mmol/L fructose. Then, tris-based extender was mixed with 200 ml fresh egg yolk and the distilled water to make 1 L extender^48^. At each generation, eight tris-based extenders were prepared by dissolving prescribed concentrations of ingredients that are presented in Table 1. For four bulls per generation, diluted semen straws were prepared in triplicate for each of the eight algorithmically generated media extenders and the control extender. To do so, diluted semen samples were cooled to 4 °C for a 4 h equilibration time in each medium with one-step processing, packaged in 0.25 ml French straws, and then frozen in a controlled rate freezer (Digitcool 007262, IMV, France) as follows: from 4 °C to −12 °C at −4 °C/min, from −12 °C to −40 °C at −40 °C/min, and from −40 °C to −140 °C at −50 °C/min before plunging into liquid N2. After at least 24 h of storage, frozen straws were thawed by transferring directly to a 37 °C water bath for 45 s before evaluation.

### 5.4 Algorithm design

A machine learning algorithm was implemented for both modeling the outcome as a function of all inputs via an additional neural network and Gaussian process regression model and the differential evolution algorithm that informs the next component selection.

The differential evolution algorithm used in this study was developed from strategy 1 (DE/random/1/bin, which has better accuracy where sharp changes may appear in the optimization space) by Storn and Price^21^ and was coded in python and output information about the test population was used for empirical testing. A schematic of the process is outlined in Figure 1. Briefly, the differential evolution algorithm randomly generates an initial population (generation 0) that spans the entire parameter space. Then, a test population (Generation 0.5) is generated from generation 0, by crossover, specific design variable values from generation 0 is used in test population or mutation which using strategy 1. Strategy 1 generates test population values by randomly selecting values from three other extenders from generation 0, we then take the difference between two of the three values and multiply by a mutation factor, then we add the result to the third value. This process is repeated for every individual in generation 0. Then, an experiment is performed with the initial population and test population and every individual is scored. Then, only the highest scoring individuals from both populations are selected. The selected individuals are then labeled generation 1. The process is then repeated, using generation 1. After 10 generations, strategy 1 was modified to increase variation in the test population. On top of strategy 1, we added the parameters’ reference value (Table 1) multiplied by a random value between ± (0-10) %. We set the rate at which crossover occurred to 0.5 (i.e., 50% of the time) and the mutation factor for strategy 1 at 0.9, these values were fell within the optimal range described by Pi et. al.^19^ for strategy 1. The algorithm was completed, and convergence was achieved when post-thaw motility between multiple generations began to plateau.

The optimal theoretical post-thaw motility model used in this study was created in Python. An artificial neural network and Gaussian process regression were created in Python using TensorFlow and Scikit-Learn module respectively. The loss function for both models were mean squared error of predicted vs measured post-thaw motility. The artificial neural network had 12 input neurons (e.g., design variables), 1 hidden layer with 12 neurons, and 1 output neuron (e.g., post-thaw motility). We used the rectified linear unit function as the activation function and to prevent over fitting we used L2 regularization value of 0.0001. After 5 generations of differential evolution, we used 10-fold cross validation to measure the theoretical fitness of each machine learning model. The 10-fold cross validation method produced 10 mean squared error estimates which are averaged. A smaller average mean squared error indicates better model fitness. Then, we trained each model using data from generation 1-5 (200 post-thaw motility – extender pairs) and predicted post-thaw motility for extenders produced by differential evolution generation 6 (32 post-thaw motility – extender pairs). The Kruskal-Wallis test was used to compare the post-thaw motility predictions from each model to experimental post-thaw motility. A p<0.05 was deemed significant. We selected the model that most closely emulated the experimental post-thaw motility as the optimal theoretical model and coupled that with differential evolution at generation 7.

### 5.5 Assessment of semen parameters post-thaw

For the first 17 generations, we used post-thaw motility as the optimization metric for our model to get our model within a comparable range to our control extender medium post thaw recovery. At generation 18, we changed to relative total motility as our optimization metric to give more power to our post thaw metrics by controlling for individual bull variability. The relative total motility was calculated by taking a ratio of algorithmic total motility over control total motility of each sample. Three additional iterations were performed using the new input parameter.

Motility was evaluated using the Sperm Class Analyzer (Microptic, Spain) capturing videos at 50 frames per second during one second^48^. The diluted semen (2.5 μl) was placed on a pre-warmed chamber slide (37 °C, Life optic slide, 20 μM depth chamber) and motility characteristics were determined using a phase-contrast microscope (Nikon, Canada) with a 10× objective at 33 °C.

### 5.6 Validation of top performing media recipes

We used a stepwise approach to identify the top algorithm-driven extenders by comparing the various motion characteristics and flow cytometry parameters in post-thaw semen in the top algorithm driven extender and Semex commercial extender as control. Experiments were implemented to compare the results of the top algorithm-derived media design recipes with the commercial control extender. The validation experiment was accomplished using 15 bulls in one trial. Bulls were considered replicates. There were not any selection criteria for the bulls as they were in normal production.

Motion characteristics were measured by the SCA as briefly explained above. Flow cytometric evaluations were performed using a BD LSRII cytometer (BD Biosciences, CA, USA). Three different lasers with wavelengths of 355 nm (25 mW laser output, ultra-violet laser), 488 nm (20 mW laser output, blue laser) and 633 nm (17 mW laser output, red laser) were used for analysis of semen samples. For each analysis, 10,000 sperm were measured, and data were saved as FCS file. Propidium iodide (PI) was used to measure the membrane integrity of sperm. PI binds to the DNA of cells with the damaged membrane and display a red fluorescence after excitation with a 488 nm laser. Sperm showing red fluorescence after excitation with the 488 nm laser were calculated as sperm with non-intact membrane or percentage of membrane damage (MD). Acrosome reaction was measured using peanut (Arachis hypogea) agglutinin (PNA) protein that can bind to proteins of the acrosome. PNA is conjugated with fluorescein isothiocyanate (FITC), a molecule that excites at 488 nm and provides a green, fluorescent emission at 518 nm. Sperm showing green fluorescence after excitation with the 488 nm laser were counted as sperm with acrosome reacted (AR). Mitotracker deepRed which fluoresce red after excitation with a 633 nm laser light was used to measure the mitochondrial activity of sperm. Mitotracker deepRed can across the plasma membrane and enter into the active mitochondria. The percentage of sperm showing red fluorescence after excitation with the 633 nm laser were counted considered as sperm with active mitochondria.

### 5.7 Statistical analysis

All statistical analysis were conducted in R (version 3.6.3) using the FSA: Fisheries Stock Analysis package^49^. To assess the predictive ability of the machine learning models, a Kruskal-Wallis test and Fligner-Killeen test were used to compare to experimental results. Then a post-hoc Dunn test was used to test pairwise relationships between all experimental results, artificial neural network post-thaw motility predictions, and Gaussian process regression post-thaw motility predictions. We also compared the variance produced by Generation 14 and 18 extenders was compared to commercial to assess optimization metric performance (64 motility – extender pairs). Generation 16 to 18 were compared using Kruskal-Wallis to assess if differential evolution was converging to an optimum. Comparisons were considered significant for p < 0.05. Medians and standard deviation were reported for all values unless otherwise specified.

## 6. Acknowledgements

Funding for this research was provided in part by the National Science and Engineering Research Council (CRD PJ531082) and Mitacs (to JB). We acknowledge that this research occurred on the traditional territory of the Anishinabewaki 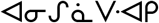, Ho-de-no-sau-nee-ga (Haudenosaunee), Omàmìwininìwag (Algonquin), Wendake-Nionwentsïo, Wabanaki (Dawnland Confederacy), N’dakina (Abenaki / Abénaquis), and Treaty 6 territory, the traditional territory of the Niitsítpiis-stahkoii 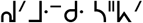 (Blackfoot / Niitsítapi 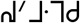), Cree, Michif Piyii (Métis), and Očhéthi Šakówiŋ. An earlier version of this work was published in the thesis of F. Tu^50^.

## 8. Contributions

FT consulted on media design, designed the ML algorithm, analyzed experimental data, contributed to writing the paper, prepared figures, reviewed and edited the paper. MB consulted on initial media components, collected experimental data, interpreted results, contributed to writing the paper, reviewed and edited the paper. PB & PV consulted on initial media components, coordinated sample collection, interpreted results, reviewed and edited the paper. MS collected experimental data, interpreted results, analyzed experimental data, prepared figures, contributed to writing the paper, reviewed and edited the paper. JB conceived of the project and developed experimental design, obtained funding, consulted on initial media components, coordinated sample collection, interpreted results, contributed to writing the paper, reviewed and edited the paper. All authors reviewed and approved the final draft.

## 9. Data Availability

The datasets generated the current study are available on request.

